# Mind the gap between functional groups and surface of magnetic nanoparticles for highly specific magnetic-based protein assays in biological medium

**DOI:** 10.64898/2026.08.29.747976

**Authors:** Rabia Amin, Yihao Wang, Florian Wolgast, Mohammad Suman Chowdhury, Nina Lehmler, Maren Schubert, Meinhard Schilling, Thilo Viereck, Aidin Lak

## Abstract

Magnetic readout-based assays are compatible with unprocessed biological samples as unbound background molecules do not interfere with magnetic signal. Yet, a true challenge is their poor specificity and susceptibility of magnetic nanoparticles (MNPs) to clusters in complex biological media, hampering their true advancement. Here, we demonstrate that the spatial organization of functional groups at the external periphery of custom magnetic nanoparticles by harnessing ultra-dense double-stranded DNA results in an efficient antibody conjugation with good accessibility toward antigen. By labeling our MNPs with anti-S protein neutralizing IgG antibody, we showcase the detection of S1 subunit of SARS-CoV-2 Spike protein in a wash-free fashion in less than five minutes in nM regime using magnetic particle spectrometer. By mixing our IgG-labelled MNPs with DMEM cell culture (10-20% FBS serum), we sense the S1 proteins in a one-pot fashion with high specificity. Our results show that by having the ultra-dense dsDNA shell on MNPs, the entropic cost of an irreversible protein binding to particle surface is high, thus allowing the formation of dynamic protein corona on the DNA shell that can be replaced with S1 protein with high affinity.

When the azide moieties are placed at the close proximity of MNPs by using non-functional dsDNA, antibody conjugation becomes inefficient, to a level not sufficient for S1 protein detection. Our study highlights the importance of spatial organization of functional moieties on the nanoscale on magnetic nanoparticles for highly specific assays in biologically complex media.

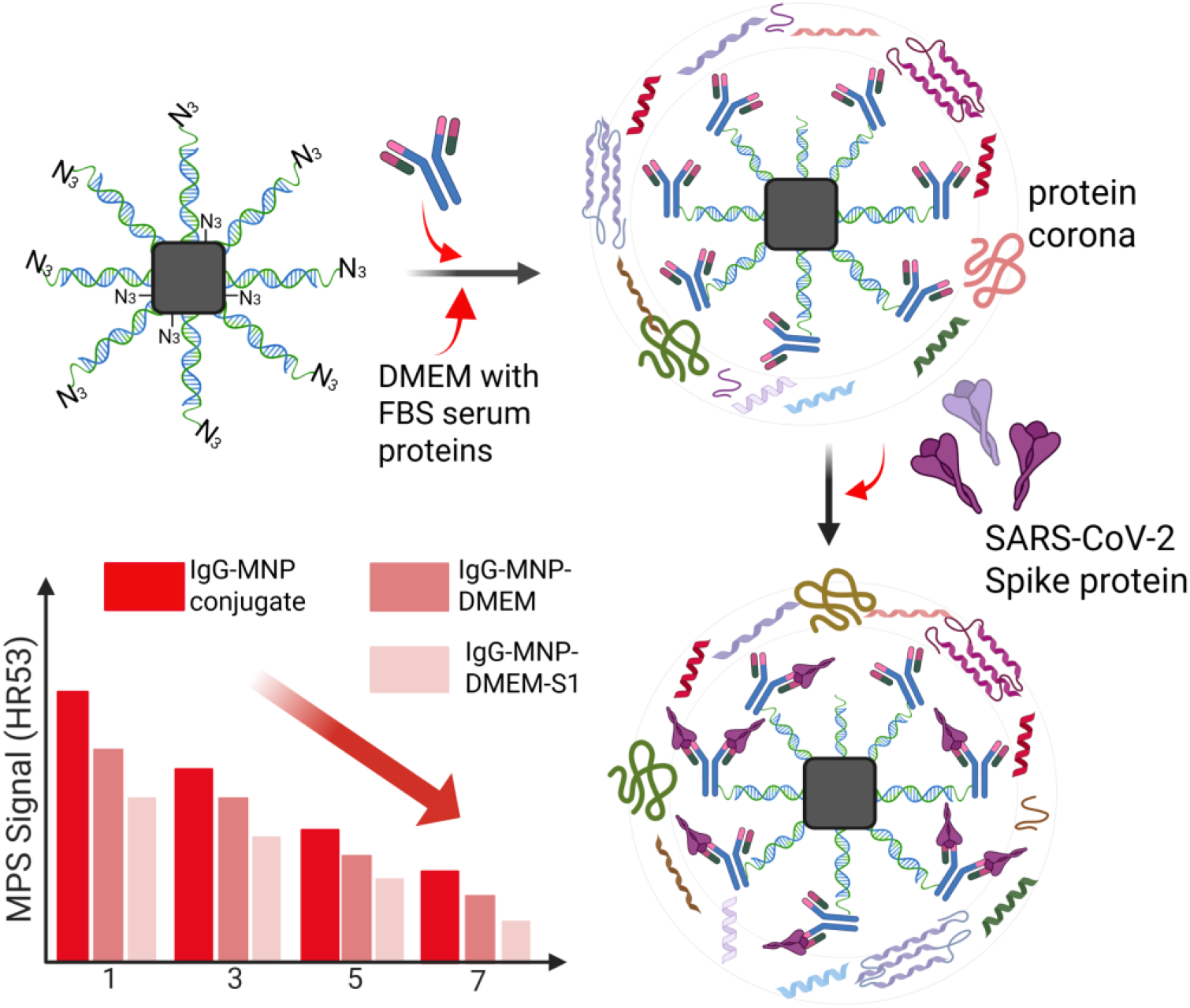

## INTRODUCTION

Nature exploits specific organization of molecules and receptors to achieve sensitive recognition with high fidelity and to recruit crucial molecules accordingly. A prime example is human T-cell receptors and their co-receptors that are assembled into certain motifs upon recognizing ligand. Over the past years, there has been some effort to endow colloidal nanoparticles with receptor spatial organization^1^, yet realizing a specific organization of functional moieties on colloidal nanoparticles remains challenging as they have non-uniform surfaces.

Colloidal magnetic nanoparticles (MNPs) offer unique features for sensing nucleic acids^2,3^ and proteins in free solution by magnetic readout. Magnetic relaxation dynamics of MNPs is highly sensitive to molecular interactions between receptors on MNPs and target in solution and can be readout by magnetic particle spectroscopy (MPS) very reliably and at ultra-low particle concentrations.^4–8^ Most magnetic bioassays are based on cross-linking MNPs upon sensing analyte.^7,8^ However, these assays are prone to false-positive, as magnetic dipole-dipole interactions between particles may lead to unspecific cross-linking. Monitoring magnetization dynamics of single MNPs by binding to proteins can resolve this issue, yet changes in particle hydrodynamic size and magnetization dynamics caused by binding 10s of kilodalton protein on MNPs is minuscule. Its realization requires a combination of high particle magnetic moment, ultra-small particle hydrodynamic size, and accessible surface receptors. Recently, it has been shown that the way a molecular binding is coupled to dynamics is not linear.^30^ If a protein binds to a receptor within an already existing surface shell on particle surfaces, the overall magnetic signal change it causes is barely detectable with MPS spectrometer. It is, therefore, crucial to increase the surface shell radially upon sensing in order to achieve highly efficient and real-time sensing scheme.

In this work, we demonstrate that by organizing functional azide moieties on the outer shell of a double-stranded DNA brush, monoclonal anti-SARS-CoV-2 antibody are labeled efficiently on MNPs that are functional against spike protein (S1 subunit) of SARS-CoV-2. In contrast, when azide moieties are buried within the brushed dsDNA layer, the antibody labeling is inefficient to an extent that no S1 protein can be detected. This works set the organization on the nanoscale as a crucial parameter to design highly specific MNP-based protein assays.

## RESULTS AND DISCUSSION

MNP-based immunoassays offers a sensitive and specific approach for detecting protein biomarkers by monitoring target-induced changes in particle hydrodynamic size. However, a key challenge in the MNP-based protein assays lies in achieving a detectable size change upon binding a target protein. Most assays rely on commercially available MNPs with core sizes ranging between 30^7,9^ to 80 nm,^8,10,11^ whereas common protein biomarkers, such as the SARS-CoV-2 spike protein (S1 subunit) and nucleocapsid protein span only 5–25 nm in size. Binding of a single protein to relatively large MNP therefore produces only a small relative increase in particle size, resulting in a negligible change in hydrodynamic size that can be difficult to detect. An equally important challenge is maintaining colloidal stability across multiple functionalization steps required to build a functional bioassay surface. We have addressed these challenges by using custom-made MNPs with core sizes below 20 nm, combining a small particle size with high magnetic moment for sensitive protein detection. We further employed a dense double-stranded DNA shell around the polymer-coated MNPs, which provides electrostatic stabilization and shields the particle surface from non-specific interactions. While the dense dsDNA shell provides enhanced colloidal stability, it may also limit the accessibility of functional groups located within the DNA brush to antibodies. Here, we show that spatial positioning of functional groups within the dsDNA shell plays a vital role in antibody conjugation and subsequent protein detection. The efficient binding of monoclonal anti SARS CoV-2 IgG antibodies (STE90-C11)^12^ to the MNP surface when azide-functionalized groups are arranged at the outer shell of the dsDNA is schematically shown in Figure 1. Conversely, no binding of IgG is seen when complementary ssDNA (comp. ssDNA) with no azide moieties hybridizes with the surface-bound ssDNA (label ssDNA).

**Figure 1.**
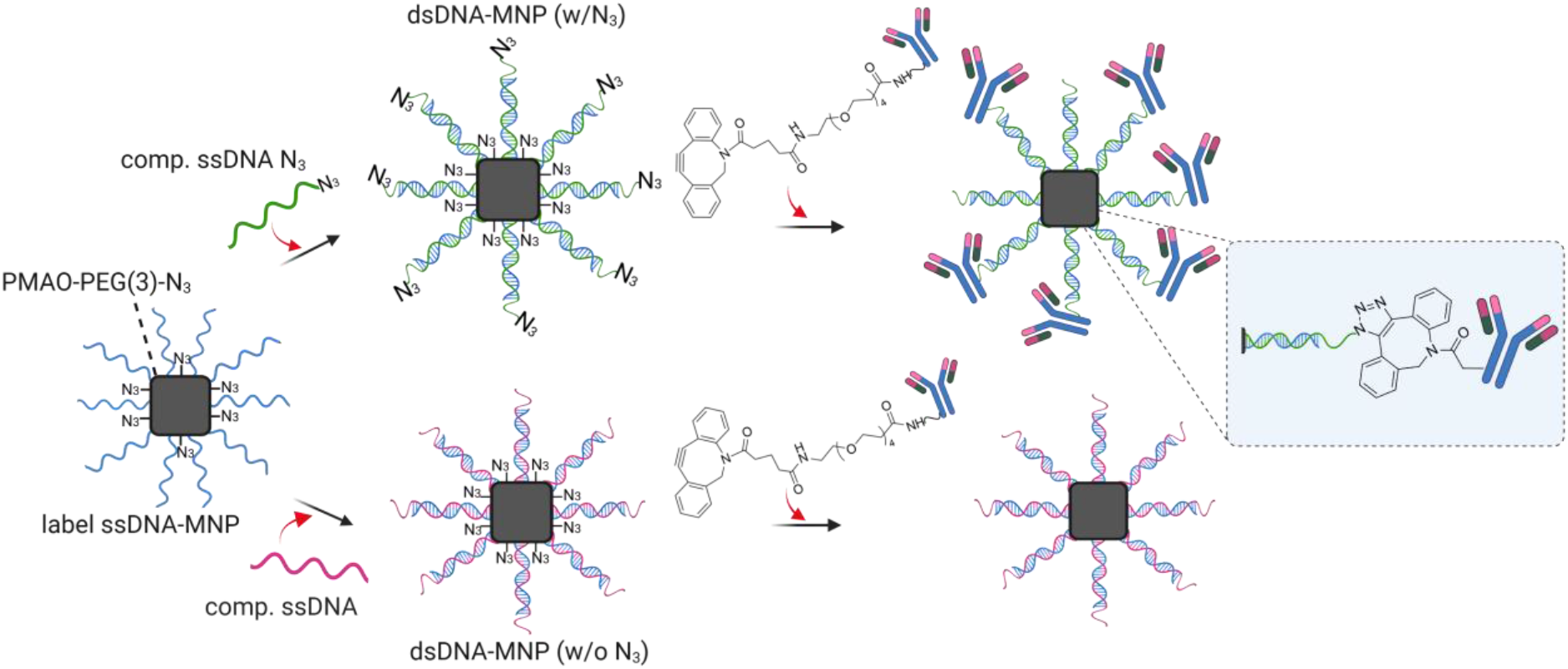
Schematic representation of the concept of minding the gap between azide functional moieties and surface of MNPs by DNA hybridization. By placing the functional groups at the extreme periphery of MNPs, monoclonal anti-SARS-CoV-2 IgG antibodies are conjugated to MNPs efficiently. In case of having azide functionality on particles surfaces, no antibodies are conjugated to MNPs.

Brownian relaxing cobalt- and zinc-doped iron oxide MNPs with cubic morphology and rounded edges were synthesized following our published protocol,^13^ with slight modifications.^14^ Transmission electron microscopy (TEM) analysis revealed a mean particle core size of 18.5 ± 2.4 nm (mean ± standard deviation) (see Supporting Information (SI), Figure S1 a,b). The MNPs were further characterized with inductively-coupled plasma optical emission spectroscopy (ICP-OES) and the stoichiometry was determined to be Co0.35Fe_2.38_Zn_0.27_O_4_. The field-dependent magnetization hysteresis loops measurements were performed using a magnetometer (MPMS Quantum Design), and yielded a maximum magnetization (*M*_max_) of 74.6 ± 4.2 emu/g (~ 114 emu per gram of Fe + Co) at 298 K (Figure S1c).

The preparation of azide-functionalized MNPs by coating with poly(maleic anhydride-*alt*-1-octadecene) (PMAO) copolymers follows our previously published protocol (see details in SI).^15^We first modified the PMAO copolymer with polyethylene glycol (PEG)(3)-azide linkers, to introduce azide moieties onto the MNP surface.^16^ Subsequently, DBCO-modified mixed ssDNA sequence (label ssDNA of 25 nucleotide, including 5 × Adenine as spacer, Table S1) was conjugated to the azide-functionalized MNPs via strain-promoted azide-alkyne cycloaddition (SPAAC) copper-free click chemistry.^15^ To have a dense DNA brush shell on MNPs, we opted for following our published protocol at high DNA grafting density (~391 DNA/MNP, see SI). Following the ssDNA labeling of MNPs, fully complementary ssDNA (comp. ssDNA of 20 nucleotides, Table S1) was added to hybridize with the surface-bound label ssDNA. To test our hypothesis on whether the spatial organization of the azide functional moieties influences the subsequent binding and accessibility to DBCO-tagged antibodies (Figure 1), we designed two sample sets. In the first set, sample was hybridized with azide-functionalized comp. ssDNA, in which azide moieties were positioned at the outer end of the DNA strand, whereas the second set was hybridized with the comp. ssDNA lacking azide groups. Subsequently, DBCO-tagged IgG were added to both sample sets to enable conjugation via the SPAAC click-chemistry approach.

To monitor how ssDNA labeling, DNA hybridization, and IgG conjugation change the particle translational and rotational dynamics, we applied a set of complementary techniques. The volume-weighted particle hydrodynamic size *D*_h_ increases from 47.8 to 75.5 and then to 83.3 nm upon DNA labeling and hybridization on MNPs, as measured by dynamic light scattering (DLS) (Table 1). Regardless of the comp. ssDNA being tagged with or without N_3_, *D*_h_ increases nearly similarly for ≈ 35 nm, showing the formation of dense brushed DNA shell on MNPs in both cases. However, after adding the IgG antibody, the particle *D*_h_ changes very differently depending on the spatial position of N_3_. In case of having N_3_ groups on the outer shell of DNA brush layer (the upper pathway of Figure 1), we observe a definite ≈ 23 nm increase in *D*_h_ upon IgG conjugation, indicating an efficient labelling of MNPs with IgG antibodies (Figure 2a and Table 1). Considering the typical dimensions of an IgG molecule with height of 8.5 nm and thickness of 4.0 nm,^17–19^ this increase is consistent with a monolayer formation of IgG in an upright (Fc-bound) orientation on the MNP surface^20–22^ (see SI for details). In contrast, when N_3_ groups are organized on the immediate surface of MNPs (the lower pathway of Figure 1), the conjugation of IgG to MNPs results in ≈ 1 nm *D*_h_ increase, implying nearly no IgG bind to MNPs. We attribute these two very different behaviors mainly to the accessibility of the azide groups depending on their spatial organization on MNPs. When they are buried within the dsDNA brush, they are sterically shielded and are therefore inaccessible to IgG.

**Table 1.**
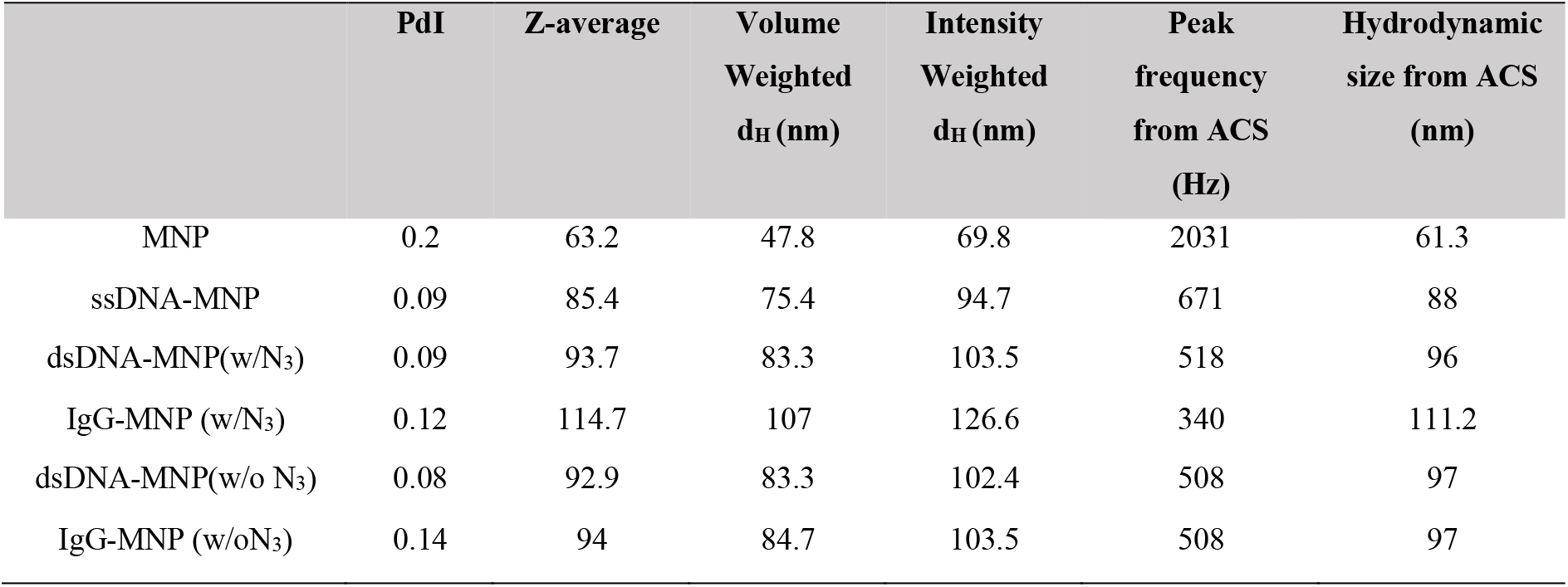
Characterization of particle hydrodynamic size and polydispersity index at each binding step, as determined by DLS, representing the Z-average, volume and intensity-weighted particle size distributions (nm). The values reported here are the average of three measurements done by DLS. Brownian relaxation peak frequencies obtained from the ACS measurements and the corresponding calculated hydrodynamic size are reported separately.

| | PdI | Z-average | Volume<br>Weighted<br>$d_H$ (nm) | Intensity<br>Weighted<br>$d_H$ (nm) | Peak<br>frequency<br>from ACS<br>(Hz) | Hydrodynamic<br>size from ACS<br>(nm) |
| --- | --- | --- | --- | --- | --- | --- |
| MNP | 0.2 | 63.2 | 47.8 | 69.8 | 2031 | 61.3 |
| ssDNA-MNP | 0.09 | 85.4 | 75.4 | 94.7 | 671 | 88 |
| dsDNA-MNP(w/N<sub>3</sub>) | 0.09 | 93.7 | 83.3 | 103.5 | 518 | 96 |
| IgG-MNP (w/N<sub>3</sub>) | 0.12 | 114.7 | 107 | 126.6 | 340 | 111.2 |
| dsDNA-MNP(w/o N<sub>3</sub>) | 0.08 | 92.9 | 83.3 | 102.4 | 508 | 97 |
| IgG-MNP (w/oN<sub>3</sub>) | 0.14 | 94 | 84.7 | 103.5 | 508 | 97 |

**Figure 2.**
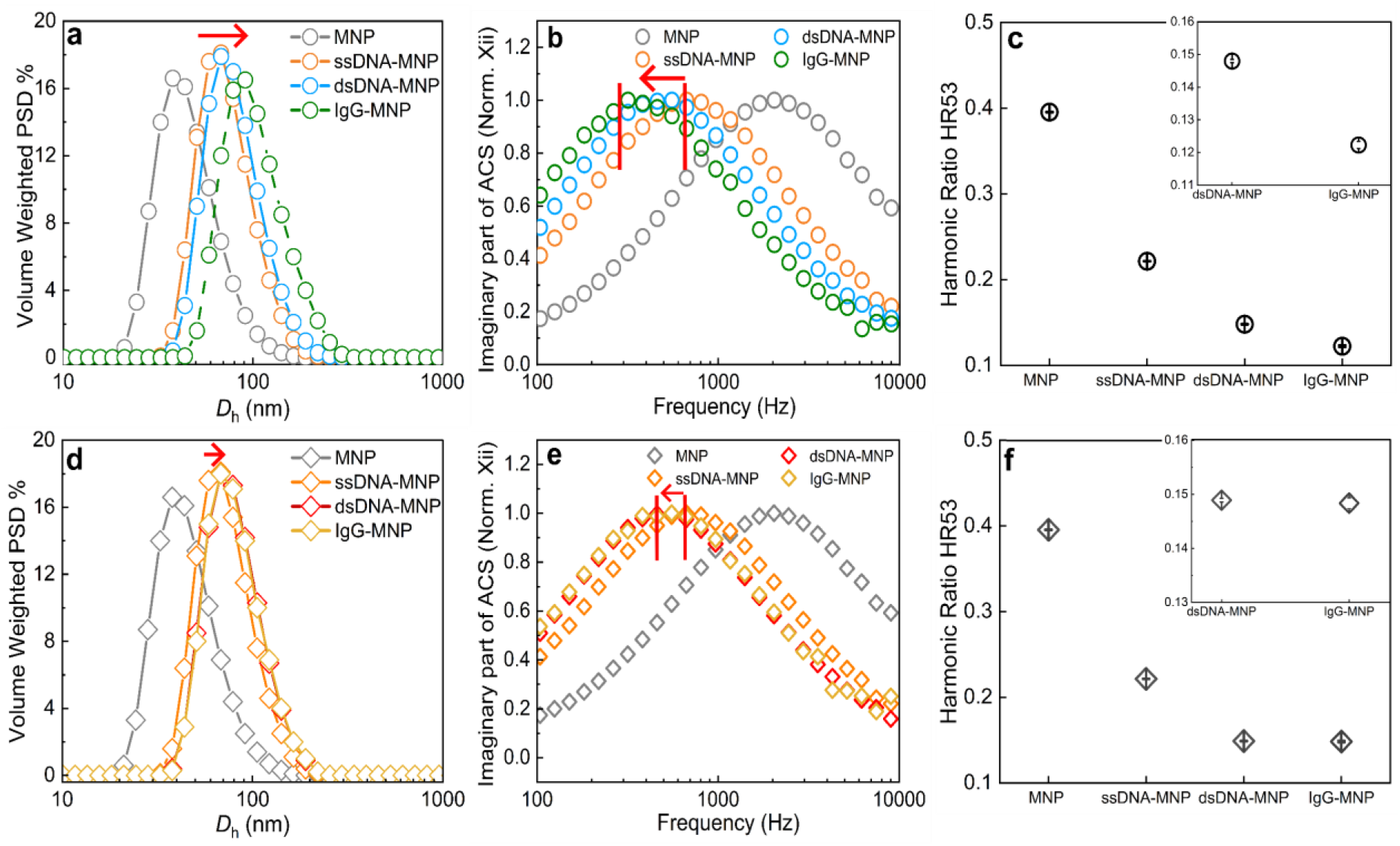
Characterization of sequential surface modifications on polymer-coated MNP from ssDNA labeling to IgG conjugation using Dynamic Light Scattering (DLS), Alternating Current Susceptometry (ACS), and Magnetic Particle Spectroscopy (MPS). Panel (a-c) show results for comp. ssDNA with N_3_ group at DNA strand terminus; panel (d-f) show results for comp. ssDNA lacking N_3_ groups. DNA hybridization is observed for both comp. ssDNA with and without N_3_ group across all three measurements. In contrast, IgG conjugation (at a fixed nominal ratio of 150 IgG/MNP) occurs exclusively in the presence of terminal N_3_ groups on the comp. ssDNA, whereas no IgG binding is observed in their absence, confirming the essential role of accessibility of functional group at the outer terminus of the dsDNA on the MNP surface. All DLS, ACS and MPS, measurements were performed at room temperature on particle suspensions of 50 μL, 150μL and 60 μL, respectively. The ACS measurements were performed at the excitation field amplitude of 0.2 mT and frequency ranging from 1 Hz to 9 kHz, while the MPS measurements were carried out at the field amplitude and frequency of 15 mT and 2 kHz. Polymer coated MNP, ssDNA-MNP and dsDNA-MNP samples were measured in PBS at a particle concentration of 12-14 nM, whereas IgG-MNP samples were measured at 4 nM.

Measuring the changes in *D*_h_ by looking into the rotational dynamics of MNPs at different labeling stages by alternating current susceptometry (ACS), we observe similar and consistent trends. Upon DNA labeling and hybridization on MNPs, the characteristic Brownian relaxation peak shifts toward lower frequencies (Figure 2 b,e). As the Brownian relaxation time is inversely proportional to the characteristic peak frequency, the systematic shifts seen here indicate a slowed relaxation process corresponding to a larger hydrodynamic volume at each binding step. While the ACS peak shifts further to lower frequencies upon the IgG conjugation in the case of N_3_ groups being positioned on the outermost periphery of MNPs, there is no visible shift in the case of N_3_ groups being inaccessible within the dsDNA brush layer.

Assessing the effect of the surface modifications differently, we looked into changes in the non-linear magnetic relaxation dynamics of MNPs by magnetic particle spectrometer (MPS) at different binding steps. Wee use the ratio of 5^th^ to 3^rd^ harmonics (HR53), since the MPS harmonics amplitudes are affected by the particle concentration, Overall, we observe behaviors consistent with the trends discussed above. Up to DNA hybridization with the comp. sequence, the HR53 drops step by step and consistently for both types of the comp. sequences (Figure 2c,f). Similar to what we observed in ACS and DLS results, while the HR53 drops further specifically upon the conjugation of IgG to N_3_ groups on the outermost periphery of MNPs, it changes marginally by having N_3_ groups inaccessible within the dsDNA brush layer. Notably, across each binding step, the peak shifts in both DLS and ACS measurements are homogenous with no visible change in the peak shape, indicating monodispersity and stability of MNPs throughout the sequential functionalization steps, as both measurements are highly sensitive to any changes in the colloidal state of MNPs such as aggregation. Converting the ACS peak frequency to *D*_h_ using Eq.9 (details in SI), these values match closely with the DLS Z-average values (Table 1).

### MNPs with antibodies at the outermost periphery are able to sense S1 subunit of SARS-CoV-2 spike protein via wash-free workflow

Establishing a wash-free protein assay demands a magnetic reader that is not falsified by diamagnetic contributions of biological matters and water. MPS spectroscopy is based on the nonlinear magnetic response of MNPs to alternating magnetic field^7,8,23,24^, making it an ideal technique, as linear magnetic contributions from background components do not affect the output signal^25^. MPS-based assays rely on Brownian relaxation of MNPs, which depends on particle hydrodynamic size. Upon molecular binding between MNPs and the target, the particle hydrodynamic volume increases, resulting in slower Brownian relaxation times and a decrease in the amplitude of the higher harmonics^7^. Higher-order harmonics are more sensitive to changes in particle relaxation dynamics than the fundamental excitation frequency. In particular, the 5^th^ harmonic drops faster than the 3^rd^ harmonic.^8,10,11^ Since, the MPS harmonics amplitudes are affected by the particle concentration, as well as changes in the relaxation times, we use the ratio of 5^th^ to 3^rd^ harmonics (HR53) for the evaluation of our assays instead of individual harmonics. The binding of a biomolecule to MNP results in the drop of the HR53 relative to the unbound sample.

Our findings so far clearly indicate that MNPs with IgG antibodies at the outer periphery (the upper pathway in Figure 1) are a good choice for developing specific and robust assays for the detection of the viral spike protein of SARS-CoV-2 (S1 subunit). To determine the optimal number of IgG per MNP at which a reasonable MPS response with high signal-to-noise ratio is obtained, we varied the nominal ratio of IgG per MNP by adjusting the concentration of IgG in the conjugation assays (Figure S3 and Table S3). The mass of IgG conjugated per MNP was determined indirectly by measuring the unbound IgG in supernatant^18,22^ using Qubit Protein assays, then converting it to actual number of IgG per MNP (SI Eq. 3 and Table S3), represented here as the average of three replicates (Figure 3a). We compared the actual number of IgG molecules conjugated per MNP with the nominal ratio and we observe an approximately linear relationship between the two. Although the actual no. of IgG on MNP surface differs substantially from the nominal ratio, increasing the nominal amount of IgG consistently resulted in greater IgG conjugation to the MNPs (Table S3). The highest measured antibody loading was 349 at a nominal ratio of 500. However, this value appears unrealistically high considering the available particle surface area. To achieve the maximum theoretical surface loading of 212 IgG/MNP, antibody must bind in a single edge-on orientation (further details in SI, Table S2). In reality, however, antibodies are expected to bind in a mixture of orientation, and the maximum achievable surface coverage will be well below the theoretical maximum. Assuming the likely scenario of equal probability of all three possible orientations of IgG conjugation, the estimated maximum number is 100 IgG per MNP (see SI), which agrees well with the experimentally determined loading at nominal ratio of 150. Another possible explanation for the relatively high IgG loading observed at the nominal ratio of 500 is the indirect method used to estimate antibody conjugation. The antibodies that are lost during the washing procedure (see method section in SI) are also included in the estimated amount of conjugated antibody, which may contribute to an overestimation of the actual number of IgG molecules bound per MNP. To monitor the changes in particle *D*_h_ with increasing IgG surface density, we measured the volume-weighted *D*_h_ of MNP samples with different IgG loadings and compared it with the actual number of IgG conjugated per MNP (Figure 3b, left axis). The particle *D*_h_ changed from 60 nm of dsDNA-MNP (w/N_3_) without IgG to a maximum of ~ 80 nm at the highest measured loading of 349 IgG/MNP (Table S4). HR53 measured at different IgG loadings showed a corresponding decrease with increasing IgG conjugation (Figure 3b, right axis), reflecting the inverse relationship between the two techniques. Despite the opposite signal directions, both measurement techniques reveal the gradual increase in *D*_h_ with increasing IgG loading on MNP.

**Figure 3.**
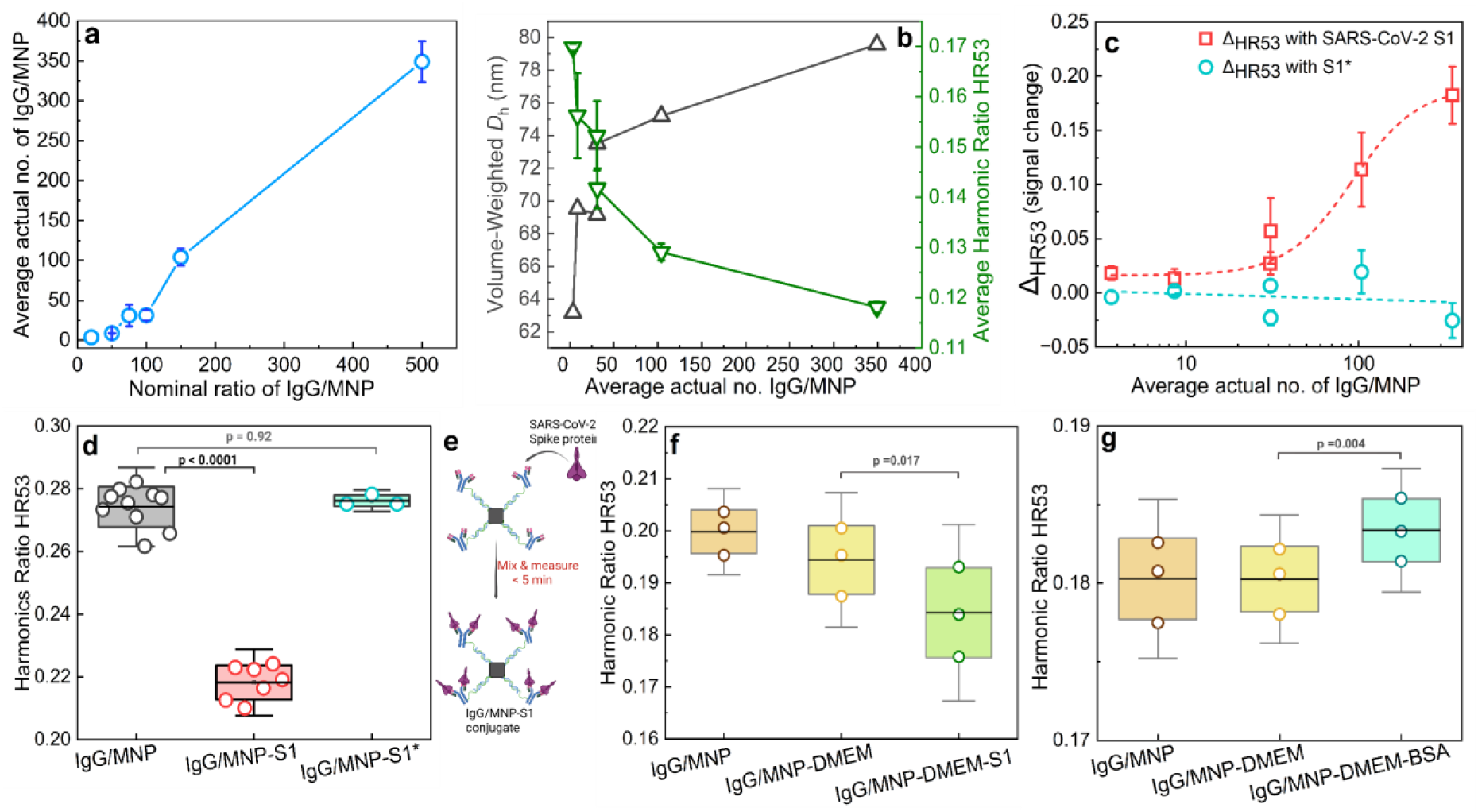
(a) Average actual number of IgG conjugated per MNP as a function of the nominal IgG/MNP ratio. Error bars represent the standard deviation between three biological replicates, prepared for each ratio from a fixed dsDNA-MNP (w/N_3_) sample at a concentration of 4nM. (b) Increase in the particle *D*_h_ with increasing IgG surface density, as reflected by the Volume-weighted *D*_h_ measured by DLS (black triangles, left axis) with a corresponding decrease in HR53 (green triangles, right axis) as a function of average actual no. of IgG/MNP. Error bars in HR53 data represents the standard deviation from three biological replicates. (c) Δ_HR53_ as a function of the average actual number of IgG per MNP upon addition of SARS-CoV-2 S1 protein (the dashed red line represent the fit of Hill equation with R^2^=0.979) and negative control S1* (the dashed green line represents the linear fit with R^2^=0.046). Error bars represent the propagated measurement uncertainties for both Δ_HR53_ with S1 and S1* (Eq 6 in SI). (d) Validation of the magnetic assay for the detection of SARS-CoV-2 S1 protein at a fixed nominal IgG/MNP ratio of 150. The p-values were calculated based on the paired two-tailed t-test. The error bars are derived from the means within the sample group. (e) Schematic of the assay workflow. (f, g) Validation of assay specificity and sensitivity in the presence of DMEM containing 10-20% FBS for detection of SARS-CoV-2 S1 (f) and control protein BSA (g). Three biological replicates of IgG/MNP samples were prepared from dsDNA-MNP (w/N_3_), at a fixed nominal ratio of 150 IgG/MNP for both data sets. S1 protein was added at a 3.6-fold molar excess relative to the IgG amount in the sample The p-values were calculated based on the paired two-tailed t-test. The error bars are derived from the means within the sample group.

Next, we added the SARS-CoV-2 Spike protein (target antigen, S1 subunit with molecular weight of 74400 g/mol) and the mutant negative control (S1* with molecular weight of 76348 g/mol), at a ratio of 1:1 of IgG:S1 (or S1*), to our IgG-MNP samples prepared at different ratios of IgG. To investigate the sensitivity of our assays, we calculated the Δ_HR53_, which corresponds to the difference between the measured HR53 of the IgG-MNP control and the sample after addition of S1 or S1*,normalized by the HR53 of the control sample (SI Eq.5). Note that the control sample here refers to the IgG-MNP at different IgG ratios. The Δ_HR53_ is shown as a function of the actual number of IgG conjugated per MNP, upon the addition of S1 and S1* protein (Figure 3c). We observe an increase in Δ_HR53_ with increasing actual number of IgG/MNP upon sensing S1, following a sigmoidal curve well described by a Hill-like equation (R^2^ = 0.979, n = 2.28, the Hill equation is given in the SI Eq. 10). This indicates that greater IgG loading on the MNP surface leads to increased S1 binding, slowing Brownian relaxation and reducing HR53. The half-maximum signal is reached at k = 91.6 ± 28.1 IgG/MNP, meaning that approximately 92 IgG per MNP are required to produce 50% of the maximum MPS response upon S1 binding. The positive Δ_HR53_ observed upon S1 addition at nominal 500 IgG/MNP ratio (Figure 3a) confirms that functional IgG are indeed present on the MNP surface at this loading condition. While the mass balance estimate of 349 IgG/MNP is likely an overestimate, the actual IgG loading at nominal 500 is greater than that achieved at nominal 150 ratio. In contrast, the addition of S1* produced no significant change in the Δ_HR53_ across all the IgG/MNP ratios, with a linear fit yielding a slope statistically indistinguishable from zero (slope = −0.005 ± 0.011, R^2^ = 0.046), confirming that the MPS signal response is specific to S1 binding and is not produced by non-specific interactions.

Using the developed workflow, we performed the magnetic assay with a fixed actual IgG/MNP ratio of 104 and a 1:1 molar ratio of IgG:S1 (or S1*), including ten samples (Figure 3d) to assess the robustness of the assay. The box plot show the distribution of Δ_HR53_ for the sample set, with the box representing mean ± 1 SD and the error bars presenting mean ± 1.96 SD. S1 protein was added to seven of the ten IgG-MNP (with dsDNA w/N_3_), and S1* was added to the remaining three. The assay readout HR53 decreases upon addition of the S1 protein whereas addition of S1* produces no significant HR53 signal change. To statistically evaluate the assay response, we performed the paired two-tailed t-test on the HR53 to see if there is a significant difference before and after addition of the S1 (or S1*) to IgG-MNP sample. The HR53 values of the IgG-MNP-S1 group were found to be significantly different from those of the IgG-MNP control with p < 0.0001, confirming that S1 binding produces a statistically significant decrease in HR53. In contrast, no significant difference was observed between the IgG-MNP-S1* group and its paired control with p = 0.92, consistent with the absence of specific binding of S1* to the immobilized antibody (Figure 3d. and Figure S4 for results on individual samples).

### MNPs with antibodies displayed at the outermost periphery retain S1 sensing in the presence of a protein corona

When nanoparticles are introduced in the biological fluids, proteins from the surrounding environment can adsorb non-specifically onto the nanoparticle surface, forming a protein corona.^26–29^ The adsorbed layer of off-target proteins can potentially mask antibody binding sites and compromise assay specificity. It is therefore critical to evaluate the performance of our IgG labeled MNPs under physiological conditions. To assess this, we performed the magnetic assay in DMEM (Dulbecco Modified Eagle’s Medium) supplemented with 10-20% FBS (fetal bovine serum) (Figure 3f,g), where off-target proteins are expected to adsorb onto the MNP surface and form a protein corona. The addition of DMEM to IgG/MNP resulted in an increase in *D*_h_ size from 68 nm for IgG/MNP to 73.6 nm for IgG/MNP-DMEM (Figure S6 and Table S5), corresponding to a slight decrease in HR53 (Figure 3f,g). This size increase is attributed to the formation of protein corona around the MNP surface.^28^ An initial experiment at 1:1 molar ratio of IgG:S1(Figure S7) showed no observable HR53 signal change, likely due to reduced accessibility of IgG binding site within the protein corona, variability in the actual number of IgG conjugated per MNP, and insufficient amount of S1 protein to compete effectively with the adsorbed off-target proteins. We therefore, repeated the experiment with 3.6-fold higher amount of S1, and observed a significant signal drop in HR53 relative to IgG-MNP-DMEM sample. The box plots show the distribution of HR53 for each condition, with the box representing the mean ± 1 SD and the error bars presenting mean ± 1.96 SD (Figure 3f,g). A paired two-tailed t-test confirmed that the HR53 values of IgG-MNP-DMEM-S1 were statistically different from those of the IgG-MNP-DMEM with p = 0.017. This demonstrates that S1 protein successfully displaces the off-target protein corona to bind the immobilized IgG on the MNP surface.

We further confirmed the assay specificity by using bovine serum albumin (BSA) as a negative control protein under the same conditions (Figure 3g). We observed a similar behaviour of slight decrease in HR53 with the addition of DMEM to IgG-MNP sample. Upon addition of BSA to IgG-MNP-DMEM, we observed a slight increase in HR53. A paired two-tailed t-test showed this change as statistically significant with p = 0.004, however, the signal change is in opposite direction to that observed with S1 addition. This increase in HR53 following BSA addition is likely due to non-specific interactions of BSA with the off-target proteins adsorbed on MNP surface rather than specific antibody binding. The opposing directionality of BSA and S1 protein responses confirms the selectivity of the IgG-MNP conjugates in a complex biological media.

To sum up, here we present a one-pot, wash-free magnetic assay for the detection of SARS-CoV-2 Spike protein in nM regime in complex biological media. We show that the spatial organization of reactive azide moieties at the distal end of the dsDNA shell is a critical parameter for efficient antibody conjugation and highly sensitive target protein detection. This accessibility of functional groups enables controlled IgG loading, with increasing surface density produces a systematic increase in hydrodynamic size and a corresponding decrease in HR53. This correlation demonstrates that antibody surface density directly influences the magnetic response of MNPs. In contrast, when the dense dsDNA shell is void of functional groups at its outer terminus, IgG labelling is inefficient to a point not sufficient for S1 protein detection. The sequential functionalization of the polymer-coated MNPs with the label and comp. DNA followed by IgG conjugation, maintained monodisperse particle populations with no observable bimodal distributions in the ACS spectra. These results indicate that majority of the MNP population was successfully functionalized and remained colloidally stable throughout each binding step, even upon introduction to complex biological medium containing DMEM supplemented with serum proteins. Operating under isothermal conditions (25ºC) and without enzymatic reagents, the assay retains the ability to detect the SARS-CoV-2 Spike protein with high specificity in the presence of off-target proteins. Importantly, the MNPs maintained a controlled hydrodynamic size throughout the functionalization and S1 binding, preserving their magnetic response for reliable detection. While the current assay enables sensitive protein detection, the limit of detection can be improved by further optimizing IgG loading density on the MNP surface. Increasing the number of accessible IgG molecules may further enhance the target binding and thus enable protein detection at lower concentrations. Owing to the simple wash-free design, it can be readily adapted for multiplexed detection of multiple protein targets and adapted to clinical metrices such as plasma present natural step towards its applicability across diagnostic settings.

## Supporting information

Supplementary information for Mind the gap between functional groups and surface of magnetic nanoparticles for highly specific magnetic-based protein

## ASSOCIATED CONTENT

### Supporting Information

The following files are given in the supporting information and are available free of charge.

Experimental details including characterization techniques, materials and methods (polymer coating procedure of MNPs, MNPs purification methods, DNA hybridization protocols, DBCO-labeling of antibodies, MNP-antibody conjugation protocol, and detection of SARS-CoV-2 Spike protein). Theoretical estimation of monolayer surface coverage of IgG, determination of actual number of IgG conjugated to MNP, size characterization of IgG-MNP conjugates, BSA specificity control for magnetic assays and magnetic assay protocol in biological medium.

## AUTHOR INFORMATION

### Corresponding Author

*Aidin Lak

### Author Contributions

A.L. conceived the research idea. R.A. designed the research, synthesized particles, performed DNA functionalization, hybridization assays, DBCO labeling of IgG, protein detection assays, DLS, ACS and MPS measurements, analyzed the data, prepared the illustrations, and wrote the first draft of the manuscript. Y.W. performed polymer coating, purification of MNPs, and STEM measurement. F.W. and T.V. designed and built the MPS spectrometer. M.S.C contributed to initial assay protocol development discussions. N.L. and M.S. designed IgG and SARS-CoV-2 Spike proteins. M.S. provided resources. A.L. designed the research idea, performed TEM imaging, analyzed data, supervised the study, and wrote the manuscript.

### Notes

The authors declare no competing financial interest.

## ACKNOWLEDGEMENTS

This work is supported by the German Research Foundation (DFG 503667663 and LA 4923/3-1), and Junior Research Group “Metrology4life”.

