## Supplementary information for Mind the gap between functional groups and surface of magnetic nanoparticles for highly specific magnetic-based protein for "Mind the gap between functional groups and surface of magnetic nanoparticles for highly specific magnetic-based protein assays in biological medium"

### Materials and Methods

#### Chemicals

Iron (III) acetylacetonate ($Fe(acac)_3$, 99.9% trace metal basis), cobalt (II) acetylacetonate ($Co(acac)_2$, ≥ 99.0%), dibenzyl ether (DBE, ≥ 98%), 1-Octadecene (ODE, 90%, and 95% pure), oleic acid (OA, 90%), poly(maleic anhydride-alt-1-octadecene) PMAO (Mn = 30,000-50,000), $NH_2$-PEG(3)-Azide, Tris-Base, EDTA, NaCl, DBCO-dPEG4-TFP ester were purchased from Sigma-Aldrich, Germany. Sodium oleate (NaOL, > 97%) was purchased from TCI, America. Zinc (II) acetylacetonate ($Zn(acac)_2$, 95%) was purchased from Merck. Ethanol, methanol, isopropanol, acetone, chloroform and bovine serum albumin (BSA) with the highest purity grade were purchased from Carl Roth, Germany. All ssDNA oligonucleotides were obtained from Biomers GmbH, Germany (Table S1). Dulbecco's Modified Eagle Medium (DMEM) supplemented with fetal bovine serum (FBS) was purchased from Thermo Fisher Scientific DE. All other chemicals and reagents were purchased from Sigma-Aldrich unless otherwise stated. No further purification of the chemicals was carried out prior to their use. Anti-SARS-CoV-2 monoclonal IgG antibody (STE90-C11), and the negative control for SARS Co-V-2 Spike protein with specific point mutations (K417N-E484K-N501Y), referred here as S1* used for magnetic assays were synthesized following our published protocol.[1] The target SARS-Co-V2 Spike protein (S1 subunit, His Tag, Batch No. 40150-V08B1, 100 μg) was purchased from Sino Biological GmbH.

#### Magnetic nanoparticle synthesis

Cobalt- and zinc-doped ferrites nanoparticles (CFZ-MNPs) were synthesized via thermal decomposition of cobalt (II) acetylacetonate, iron (III) acetylacetonate and zinc (II) acetylacetonate in a mixture of 1-Octadecene, dibenzyl ether, oleic acid and sodium oleate at 290ºC for 30 min following our previously published protocol.[2,3]

#### Modification of poly(maleic anhydride-alt-1-octadecene) with $NH_2$-PEG (3)-azide linker

PMAO was modified with $NH_2$-PEG (3)-azide linker via anhydride ring opening reaction[4] following our published protocol.[5]

#### Polymer coating of MNPs

The polymer coating of CFZ-MNPs was performed following the protocols of Pellegrino et al.[6] and Lak et al.[5] with slight modifications. In a typical procedure, 1 mL of particle suspension in chloroform (4.43 g(MNP)/l) and 4 ml of PMAO-PEG polymer solution (~ 35 mg/ml), corresponding to 500 polymer units per $nm^2$ of a MNP, were further diluted with 9 and 6 mL of chloroform, respectively, in a 30 mL glass vial and sonicated for 10 min. Then, during sonication, the particle suspension was slowly added dropwise to PMAO-PEG solution, and kept sonicating for another 10-15 min to get a homogenous solution. After mixing, the vial was left on a shaker at room temperature overnight at 500 rpm. Next, the mixture was poured into a 250 ml round bottom flask, and connected to a rotary evaporator. Subsequently, chloroform was removed over 5-6 h through a stepwise reduction of pressure from 980 mbar to a final value of ~ 350 mbar and gradual increase of temperature from 29ºC to 34ºC. After complete removal of chloroform, particles were resuspended in 10 mL of sodium borate buffer (pH 8.7) by sonication for 1 h at ~ 45ºC. The particle suspension was then concentrated to 1 mL using spin filtration (Amicon regenerated cellulose spin filter, 15 ml, 30 KDa cutoff size) at 3200 rpm and 20ºC for 30 min.

**Purification and particle fractionation**

To remove unreacted polymer and polymeric micelles that are formed during the polymer coating process, the particles were fractionated by centrifugation on non-continuous sucrose gradient (10%: 40%: 60%, from top to bottom, each fraction 4 mL) columns, following our previously published protocol.[5] Typically, 500 µL of PMAO-PEG coated particle suspensions in borate buffer were loaded into the sucrose centrifuge columns and centrifuged for 90 min at 4500 rpm and 4ºC. The single polymer coated nanoparticles were collected in the upper 10% sucrose band using a long needle. The remaining sucrose was then removed through 3 rounds of Amicon filtration (Amicon spin filter, 15 mL, 50 kDa cutoff size) at 3500 rpm and 20℃ for 13 min. The particles were thoroughly resuspended in borate buffer after each centrifugation round by pipetting and shaking. Finally, the particles were upconcentrated to 1 mL in borate buffer. To further purify the particles of any remaining polymer micelles, one round of magnet washing was performed as previously described[5]. A 1.5 mL MACS column (Miltenyi Biotec) in combination with a MiniMACS separator (permanent magnet) was used. Finally, the particles were dispersed in 800 µL of TE buffer (5 mM Tris-base, 5 mM NaCl, 1mM EDTA, pH 7.3) and stored at room temperature for further use.

**Particle size fractionation using agarose gel electrophoresis**

To isolate the fraction of MNPs that are singly encapsulated within a PMAO layer, native agarose gel electrophoresis (AGE) was performed. A 1% agarose gel was prepared by dissolving 0.5 g agarose in 50 mL of 1× TAE (Tris, Acetate, EDTA) buffer using microwave irradiation for 2 min. After cooling for 5 min, the homogeneous gel solution was poured into a gel mold and allowed to solidify for 30 min. The cast gel was then placed in an electrophoresis chamber from Bio-Rad Laboratories, and the samples mixed with 20% sugar solution were loaded into the wells. Electrophoresis was carried out in 1× TAE running buffer at 90 mA for 30 min to fractionate different populations depending on their size and charge. Subsequently, the most migrating particle band (3–4 mm wide), corresponding to single polymer-coated MNPs, was excised from the gel. Finally, the particles were collected by squeezing the gel fragments using parafilm covered glass slide.

**Labeling of ssDNA to MNPs**

We functionalized polymer coated MNPs with mixed-sequence label ssDNA strands following our published protocol[5] with slight modifications. In a typical experiment, 800 μL of polymer coated MNP suspension in TE buffer (≈ 16 nM) was mixed with 493.5 μL of label ssDNA in TE buffer (100 μM) to obtain a ssDNA nominal grafting density of 1.2 ssDNA/nm$^2$ of MNP. Each ingredient was gently vortexed before pipetting to ensure homogenous sample preparation. After mixing, the sample was vortexed and sonicated for 20 s and incubated overnight at room temperature. To maximize the binding efficiency of ssDNA to MNPs, a salt aging process was applied the following day using 5 M NaCl. We increased NaCl concentration in the mixture incrementally by 100 mM every hour up to 400 mM. After every salt adjustment step, the mixture was homogenized by vortexing and sonicating for 20 s. At the end of the salt aging process, the mixture was incubated at room temperature overnight. Next, the excess of label ssDNA was washed out by two rounds of centrifugation at 13000 rcf and 10ºC for 12 min. Finally, the sample volume was adjusted to 800 μL to obtain the original particle concentration. The sample was stored at 4ºC until further use.

**Hybridizing complementary ssDNA to ssDNA-labeled MNPs**

To the label ssDNA functionalized MNP sample, we then added fully complementary ssDNA (comp. ssDNA with $N_3$ groups at 5' end and without $N_3$ groups at 5') at 150 mM NaCl in TE buffer (pH 7.3). The comp. ssDNA was added at an excess ratio of $R_{comp.DNA/label\ DNA} = 2:1$. The reaction mixtures were then incubated for 24 h in an Eppendorf thermomixer at 25ºC with 10 min intervals, alternating between 5 min shaking at 550 rpm and 5 min at 0 rpm. The excess ratio ensures hybridization of all available label ssDNA grafted on MNPs, facilitating the maximal antibody grafting in the following steps. Next, the DNA hybridized MNP sample was transferred into PBS buffer (pH 7.4) by one round of centrifugation at 13000 rcf and 10ºC for 12 min, through which unreacted comp. ssDNA are also removed. The samples (here referred as dsDNA-MNP w/$N_3$ and dsDNA-MNP w/o $N_3$) were stored at 4ºC until further use.

**Table S1.** DNA sequences used in this work for DNA hybridization protocol.

| Name | Length | Sequence | GC content | $T_m$ |
|---|---|---|---|---|
| Label ssDNA | 20 nt | DBCO 5´-(AAA AA) A GGA TTG CGG GTG CCA ATG T-3´ | 44.0 % | 64 ºC |
| Comp. ssDNA (w/$N_3$) | 20 nt | Azide 5´-ACA TTG GCA CCC GCA ATC CT-3´ | 55.0 % | 59 ºC |
| Comp. ssDNA (w/o $N_3$) | 20 nt | 5´-ACA TTG GCA CCC GCA ATC CT-3´ | 55.0 % | 59 ºC |

### DBCO-labeling of Antibodies

We functionalized the monoclonal IgG antibody (STE90-C11) in PBS buffer (pH 7.4) with DBCO-dPEG4-TFP ester through nucleophilic reaction of the TFP ester with primary amines on lysine residues of the IgG, yielding DBCO-functionalized IgG. We used the following equation to calculate the volume of DBCO-dPEG4-TFP ester required to functionalize the IgG.

$$V_{DBCO-dPEG4-TFP\ ester} = \frac{\frac{IgG_C}{IgG_{MW}} \times V_{IgG}\ \times arbitrary\ \#\ of\ lysine}{DBCO_{stock\ sol}\ \ \times arbitrary\ \#\ of\ DBCO\ per\ IgG} \qquad \text{Eq.1}$$

With $IgG_c$, $IgG_{MW}$ and $V_{IgG}$ is the concentration, molecular weight (150 kDa), and volume (µL) of the IgG. One thing of note here is that the number of lysine residues per IgG varies, therefore, an arbitrary value of 92 lysine residues per IgG was taken for calculations. To estimate the number of DBCO per IgG, the number of lysine residues was divided by an arbitrary factor of 12.

In a typical functionalization protocol, 200 µL of IgG (1.39 mg/mL, 150 kDa) was mixed with 4.05 µL of DBCO-dPEG4-TFP ester, at a concentration of 3.5 mM in a 0.5 mL low-protein bind Eppendorf tube. After mixing, the tube was vortexed immediately at power 3 for 60 s, using 10 s pulses, with brief pauses between each pulse until a total of 6 pulses were completed. Mixing by pipetting was avoided. The reaction mixture was incubated at 25ºC for 2 h, followed by overnight incubation at 4ºC, without shaking. Free DBCO was removed using Zeba Spin Desalting Column (7 kDa MWCO) the following day. The concentration of IgG and DBCO were measured with UV-Vis absorption spectroscopy at 280 nm for the IgG as 11 µM and at

309 nm for DBCO as 53.3 μM. The number of DBCO molecules per IgG was calculate as 4.8 by dividing the molar concentration of DBCO by IgG.

**Click reaction between DBCO-labeled IgG and dsDNA-MNP**

In a typical protocol, 90 μL of 4 nM dsDNA-MNP sample was mixed with DBCO-labeled IgG at a concentration of 11μM in a 0.5 mL low-protein bind Eppendorf tube. After mixing, the tube was vortexed immediately at power 3 for 60 s, using 10 s pulses, with brief pauses between each pulse until a total of 6 pulses were completed. Mixing by pipetting was avoided. The reaction mixture was incubated at 4ºC for 16-17 h, followed by 4 h incubation at 25ºC, without shaking. Next, excess IgG was washed out by two rounds of centrifugation at 14000 rcf and 12ºC for 15 min. The following equation was used to calculate the volume of IgG ($V_{IgG}$) used for functionalizing dsDNA-MNP.

$$V_{IgG} = \frac{n_{dsDNA-MNP} \times V_{dsDNA-MNP} \times ratio\ of\ IgG\ per\ MNP}{IgG_c} \qquad \text{Eq. 2}$$

where, $n_{dsDNA-MNP}$ and $V_{dsDNA-MNP}$ are the particle concentration (nM) and volume of dsDNA-MNP, and $IgG_c$ is the concentration of IgG as calculated by the Beer-Lamber law using the absorption measured by the UV-Vis spectroscopy.

**Theoretical estimation of monolayer surface coverage of IgG according to IgG footprint**

The number of IgG molecules forming a monolayer on a cubic MNP was estimated knowing the footprint of IgG and the surface area of MNP. The surface area of a cubic dsDNA-MNP is given as 7200 $nm^2$, considering a volume-weighted particle hydrodynamic size of 60 nm, as obtained from DLS measurements.

Given the IgG dimensions[7,8] as a = 14.5, b = 8.5, c = 4 nm, three IgG footprints are possible as shown in Figure S2 a,b and summarized in Table S2:

**Table S2.** Possible orientations of IgG of MNP and approximate surface coverage of MNP.

| Orientation of IgG on surface | IgG footprint ($nm^2$) | IgG/MNP |
|---|---|---|
| (A) Edge-on | $8.5 \times 4 = 34\ nm^2$ | 212 |
| (B) Fc- bound (straight upright) | $14.5 \times 4 = 58 nm^2$ | 124 |
| (C)Flat-on (lying) | $14.5 \times 8.5 = 123 nm^2$ | 58 |

Assuming equal probability for all three footprints, average (footprint) = (A+ B+ C) / 3

$$\text{IgG/MNP (theoretically)} = \frac{\text{surface area of MNP}}{footprint\ IgG} = \frac{7200}{(34+58+123)/3} = \frac{7200 \times 3}{215} = 100\ \text{IgG/MNP}$$

**Determination of actual number of IgG conjugated to MNP**

Three biological replicates of 90 µL of IgG/MNP samples were prepared from dsDNA-MNP (w/$N_3$) at a particle concentration of 4 nM, with varying nominal ratio of IgG per MNP. Following the conjugation of IgG to MNP, excess IgG was removed by centrifugation and the supernatant containing unreacted or free IgG was collected. The concentration of IgG in the supernatant were determined by the Qubit Protein Assay Kit (Thermo Fischer Scientific, Q33211) following the manufacturer's instructions. The mass of IgG conjugated to the MNP was determined by mass balance as[9,10]

$$m_{\text{IgG conjugated}} = m_{\text{IgG added}} - m_{\text{IgG in supernatant}}$$

where $m_{\text{IgG conjugated}}$ is the actual mass of IgG conjugated per MNP, $m_{\text{IgG added}}$ is the initial known mass of IgG added, and $m_{\text{IgG in supernatant}}$ is the mass of unbound IgG in the supernatant. The actual number of IgG conjugated per MNP was then calculated as

$$\text{IgG/MNP} = \frac{m_{IgG\ conjugated} \times 10^{-9}}{Mw_{IgG} \times n_{MNP}} \qquad \text{Eq. 3}$$

where $m_{\text{IgG conjugated}}$ is the actual mass of IgG conjugated per MNP , molecular weight of IgG is 150 KDa and $n_{MNP}$ are the moles of MNP.

**Table S3.** Qubit Protein Assay analysis of IgG conjugation to MNPs at different nominal ratios. The average actual number of IgG/MNP was calculated from the valid replicates only. Negative values indicate measurements exceeding the initial mass added, attributed to assay variability, '-' denotes unreliable data.

| Nominal ratio IgG/MNP | Initial mass of IgG added (ng) | IgG mass in supernatant | | | IgG mass conjugated to MNPs | | | Actual no. of IgG conjugated to MNP | | | Average actual no. of IgG/MNP |
|---|---|---|---|---|---|---|---|---|---|---|---|
| | | Replicate A (ng) | Replicate B (ng) | Replicate C (ng) | Replicate A (ng) | Replicate B (ng) | Replicate C (ng) | Replicate A (ng) | Replicate B (ng) | Replicate C (ng) | |
| 20 | 1080 | 1200 | 880 | Too low | -120 | 200 | -- | -2.2 | 3.7 | -- | 3.7 |
| 50 | 2700 | 2832 | 2224 | 2240 | -132 | 476 | 460 | -2.45 | 8.8 | 8.5 | 8.6 |
| 75 | 4050 | 3752 | 1960 | 1960 | 834 | 2090 | 2090 | 15.4 | 38.7 | 38.7 | 30.9 |
| 100 | 5400 | 3752 | 4040 | 3344 | 1648 | 1360 | 2056 | 30.5 | 25.2 | 38.0 | 31.3 |
| 150 | 8100 | 2040 | 2856 | -- | 6060 | 5244 | -- | 112.2 | 97.1 | -- | 104 |
| 500 | 27000 | 8320 | 6688 | 9440 | 18680 | 20312 | 17560 | 345.9 | 376.2 | 325.2 | 349 |

### Size characterization of IgG-MNP conjugates prepared from dsDNA-MNP (w/$N_3$)

To estimate the IgG layer formation on the outer terminus of dsDNA-MNP (w/$N_3$), samples prepared at varying IgG/MNP ratios were characterized with DLS[11,12]. The dependence of the hydrodynamic size ($D_h$) on both the nominal and actual number of IgG per MNP is shown in Table S4. As the IgG loading increases, the volume-weighted $D_h$ increases from 60 nm of dsDNA-MNP (without IgG) to a maximum of ~ 80 nm at an actual number of 349 IgG/MNP. At maximum IgG loading (actual 349) $D_h$ size increases by ~ 20 nm. As described earlier, IgG is Y-shaped protein with a height of 8.5 nm, width of 14.5 nm and thickness of 4.0 nm, thus a conjugated IgG on MNP surface is expected to increase $D_h$ by 8 nm if IgG is in a flat-on (lying) orientation, or by 17 nm if oriented perpendicular (Fc-bound, upright) to the MNP surface (Figure S2c). We observe an increase of ~ 9 nm at the inflection point[12] at an actual IgG loading of 8-9 IgG/MNP indicating that IgG are arranged in a flat-on orientation at low density. Further increase in $D_h$ at higher IgG loading towards 17-20 nm indicates that IgG would begin to take upright (Fc-bound) orientation. This suggests that as the density of IgG increases, IgG begins to reorient from a flat-on towards upright configuration, rather than forming a second layer.

**Table S4.** Volume-Weighted $D_h$ of IgG-MNP conjugates at varying nominal IgG/MNP ratios, determined by DLS measurement. The $\Delta D_h$ indicates the increase in $D_h$ relative to the dsDNA-MNP (w/$N_3$) prior to IgG conjugation ($D_H$ = 60 nm).

| Nominal ratio IgG/MNP | Average actual no. of IgG/MNP | Volume-Weighted $D_h$ nm | $\Delta D_h$ (from dsDNA-MNP(w/$N_3$)) |
|---|---|---|---|
| 20 | 3.7 | 63.2 | 3.2 |
| 50 | 8.6 | 69.5 | 9.5 |
| 75 | 30.9 | 69.2 | 9.2 |
| 100 | 31.3 | 73.5 | 13.5 |
| 150 | 104 | 75.2 | 15.2 |
| 500 | 349 | 79.6 | 19.6 |

### Binding of SARS-CoV-2 Spike protein with IgG-labelled dsDNA-MNP and measuring with MPS

The SARS-CoV-2 Spike protein (S1 subunit with molecular weight of 74400 g/mol) was added to the 60 µL of the washed IgG labeled dsDNA-MNP in a glass measurement vial, vortexed slightly and incubated for roughly 5 minutes before measuring with our custom-made 2 kHz

MPS device. To minimize the background noise, a blank sample containing PBS buffer was measured and subtracted automatically before each assay sample. Each sample was measured five times, and the average and standard deviation were calculated.

**BSA specificity control for magnetic assays**

To assess the specificity of our magnetic assays and to confirm that BSA doesn't interfere with the IgG-MNP conjugates, 0.5% BSA was added to the IgG-MNP samples. Two IgG-MNP samples of 4 nM particle concentration were prepared at a nominal IgG/MNP ratio of 150. Following IgG-MNP conjugation, excess IgG was removed by centrifugation as described earlier. A 0.5% BSA solution was prepared by adding 0.05 g BSA in 10 ml PBS. BSA was then added to each sample at a 10-fold excess relative to the IgG amount present in the samples and incubated at 25ºC, for 30 min with gentle shaking at 300 rpm. Afterwards, the free BSA was not removed, and the samples were measured directly with our custom-built MPS device. The HR53 value remained unchanged upon BSA addition compared with the IgG-MNP samples without BSA, indicating that BSA doesn't bind non-specifically to the IgG-MNP surface (Figure S5) .

**Wash-free magnetic assays in complex biological medium**

To evaluate the performance of our MNPs with IgG conjugated at the distal terminus of the dsDNA shell (dsDNA-MNP (w/$N_3$)), we performed the magnetic assay in a complex biological medium, DMEM supplemented with 10-20% FBS. A total of six IgG-MNP samples at 4 nM particle concentration were prepared at a fixed nominal IgG/MNP ratio of 150. After IgG-MNP conjugation, excess IgG was removed by centrifugation as described earlier, and 45 µL of each sample was measured with MPS in a glass measurement vial. To maintain a simple, one-vial wash-free assay, 15 µL of DMEM was added to the IgG-MNP sample in the glass vial, and the vials were then incubated at 25ºC for 30 min with gentle shaking at 300 rpm. The IgG-MNP-DMEM samples were measured again with MPS, after which the target S1 protein was added to three IgG-MNP-DMEM and control BSA protein was added to the remaining three in same glass vial (Figure 3f,g). The samples were vortexed and incubated for roughly 5 min before measuring again with our custom-built 2 kHz MPS device. The volumes of IgG-MNP (45 µL), DMEM (15 µL), and S1 (5 µL) (or BSA, 5 µL) were chosen to keep the total volume in the glass measurement vial below 65 µL, as required for MPS measurement.

The increase in the volume-weighted $D_h$ of IgG-MNP conjugates suspended in DMEM after 30 min of incubation at 25°C suggests the formation of protein corona around the MNPs. Further increase in $D_h$ following the addition of S1, when added at a 3.6-fold excess relative to IgG amount in sample, which has a high affinity for IgG, were able to displace the off-target proteins and bind to the immobilized IgG on MNP surface (Table S5 and Figure S6). In contrast, no increase in $D_h$ was observed after BSA addition; instead a $D_h$ remained comparable to that of IgG-MNP conjugate. This suggests that BSA may interact non-specifically with the off-target proteins rather than bind specifically to IgG. A consistent trend was observed in MPS measurements, where HR53 increased upon BSA addition relative to IgG-MNP-DMEM samples (Figure 3g), further supporting the absence of specific BSA binding.

**Table S5**. Characterization of particle hydrodynamic size and polydispersity index at each binding step, as determined by DLS, representing the Z-average, and volume-weighted particle size distributions (nm). The values reported here are the average of three measurements done by DLS. The S1 protein was added at a 3.6-fold molar excess relative to the IgG amount in the sample.

| | **PdI** | **Z-average** | **Volume Weighted $D_h$ (nm)** |
|---|---|---|---|
| IgG-MNP (w/N3) | 0.15 | 82.2 | 68.0 |
| IgG-MNP-DMEM | 0.27 | 79.1 | 73.6 |
| IgG-MNP-DMEM-S1 | 0.23 | 86.4 | 79.8 |
| IgG-MNP-DMEM-BSA | 0.25 | 71.6 | 65.5 |

**Analysis of the MPS-based assay readout and measurement uncertainties**

For analysis of the MPS data, the harmonic ratio between the 5th and the 3rd harmonic is calculated as

$$HR53 = \frac{H5}{H3} \qquad \text{Eq. 4}$$

The relative change in harmonic ratio $\Delta_{HR53}$ is calculated as

$$\Delta_{HR53} = 1 - \left(\frac{HR53_{sample}}{HR53_{control}}\right) \qquad \text{Eq. 5}$$

The relative change $\Delta_{HR53}$ represents the signal change attributed to the target detection. The uncertainty of $\Delta_{HR53}$ was determined by standard error propagation of the individual uncertainties of $HR53_{control}$ and $HR53_{sample}$ :

$$\sigma\Delta HR = \sqrt{(\frac{\sigma HR_{sample}}{HR_{control}})^2 + (\frac{HR_{sample} \times \sigma HR_{control}}{HR_{control}^2})^2} \quad \text{Eq. 6}$$

Where $\sigma HR_{sample}$ and $\sigma HR_{control}$ are the standard deviations of the harmonic ratio of the sample and control, respectively.

**Calculation of hydrodynamic diameter from ACS peek frequency**

The Brownian relaxation time $\tau_B$ of MNP is related to peak frequency $f$ of the imaginary part of the ACS spectrum by:

$$\tau_B = \frac{1}{2\pi f} \quad \text{Eq. 7}$$

The Brownian relaxation time $\tau_B$ is further related to the hydrodynamic size $D_h$ of MNP as

$$T_B = \frac{3\eta D_h}{k_B T} \quad \text{Eq. 8}$$

Where $\eta$ is the viscosity of medium, $k_B$ is the Boltzmann constant, and T is the temperature. MNPs exhibit irregular cubic morphology with rounded edges and are therefore approximated as spheres for hydrodynamic size calculations. The hydrodynamic volume is given as $V_H = \pi D^3/6$. The $D_h$ size is calculated as

$$D_h = (\frac{k_B T}{\pi^2 \eta f})^{1/3} \quad \text{Eq.9}$$

**Four parameter Hill Equation**

A four-parameter Hill equation was used to describe the sigmoidal dependence of the data

$$y(x) = A_1 + (A_2 - A) \times \frac{x^n}{k^n + x^n} \quad \text{Eq. 10}$$

where $A_1$ and $A_2$ are the lower and upper asymptotes of the sigmoidal responses, $k$ is the half-maximum constant at which the response reaches the midpoint between $A_1$ and $A_2$, n is the Hill coefficient describing the cooperativity and steepness of the transition, and x is the independent variable.

#### Characterization methods

#### Transmission Electron Microscopy (TEM)

Transmission electron microscopy analysis were carried out using a JEOL TEM microscope operating at an accelerating voltage of 100 kV. The samples were prepared by drop-casting 5 µL of particle suspension in chloroform on a 300 mesh Formvar-carbon-coated copper grid and allowing the solvent to completely dry inside a fume hood. TEM images were analyzed using ImageJ software. The size distribution and average particle size were calculated by measuring at least 200 nanoparticles.

#### Inductively-coupled plasma optical emission spectroscopy (ICP-OES)

ICP-OES was performed on a Varian (715ES) instrument to determine the concentration of elements of interest. Briefly, 50 µL particle suspension in chloroform and 1 mL aqua regia (HCl: $HNO_3$ at 3:1 (v/v)) was pipetted into 10 mL volumetric flask. Next, the samples were incubated at 60 ºC for 1 h to facilitate the digestion process and then left inside fume hood overnight. The next day, the samples were diluted with Mili-Q water up to the 10 mL grading level. To convert, Fe, Co, and Zn concentrations to particle number concentrations, the unit cell composition of an FCC crystal structure was used, with a lattice constant of $a$ = 8.40 Å.

#### Magnetic Property Measurement System (MPMS)

The magnetization hysteresis loops were measured using MPMS (Quantum Design). The sample was prepared by pipetting 20 µL of particle suspension in chloroform, at a particle concentration of 4.098 g/L in a designated sample holder, and allowing the solvent to evaporate completely at room temperature inside a fume hood. The hysteresis loops were recorded at 298 K in the vibrating-sample magnetometry (VSM) mode at magnetic fields between -7 and 7 T. The applied magnetic fields were corrected for a remanence field in the chamber and the magnets by measuring the palladium standard sample. The coercive fields were corrected accordingly. Measurement errors were obtained from combining the results of three independent measurements and error propagation.

#### Dynamic Light Scattering (DLS)

DLS measurements were performed using a Malvern Zetasizer instrument at 173° backscattered measurement mode. The DLS was measured at room temperature on 50 µL particle suspension. Polymer-coated MNP, ssDNA-MNP, and dsDNA-MNP samples were measured at 12–14 nM, and IgG-MNP and IgG-MNP-S1 (or S1*) samples were measured at 4 nM particle concentration.

**Complex ac-susceptibility (ACS) spectroscopy**

The ACS spectra was recorded using a custom setup with an amplitude of 0.2 mT, with varying excitation frequencies, ranging from 1 Hz to 9 kHz[13]. All the measurements were performed at room temperature on 150 µL of particle suspension at slightly varying concentration of 12-14 nM for all samples including polymer-coated MNPs, ssDNA-MNP and dsDNA-MNP in PBS buffer, and 4 nM particle concentration of IgG-MNP and IgG-MNP-S1 (or S1*).

**Determination of DNA grafting number on MNPs**

To determine the actual number of ssDNA onto MNP surface at a grafting density of 1.2, we followed our previously established protocol[5], and measured UV absorbance of the supernatants following DNA functionalization after centrifugation. For polymer-coated MNPs with a hydrodynamic diameter of 42 nm, we quantified approximately 391 grafted ssDNA strands per particle.

**Determination of DBCO loading per IgG**

To determine the number of DBCO per IgG, we performed UV-VIS absorbance on DBCO-labelled IgG, and measured absorbance at 280 nm for IgG and at 309 nm for DBCO. The concentrations of both DBCO and IgG were calculated in µM using the Beer-Lambert law gives as:

$$c = \frac{A}{\varepsilon . L} \qquad \text{Eq. 11}$$

where, c is the calculated molar concentration, A is the absorbance measured, ε is the molar extinction coefficient (12000 $M^{-1}cm^{-1}$ for DBCO and 206700 $M^{-1}cm^{-1}$ for IgG), and $L$ is the optical path length of 0.1 cm. The number of DBCO molecules per IgG was then determined by dividing the molar concentration of DBCO by that of IgG.

**Magnetic Particle Spectroscopy (MPS)**

MPS measurements on MNPs in PBS buffer were carried out on a custom-built setup, which is specifically designed for sensitive magnetic bioassays. The setup operates at an excitation frequency of 2 kHz and a magnetic field amplitude of $\mu_0 H$ = 15 mT. The measurements were performed on 60 µl particle suspensions at room temperature. Polymer-coated MNP, ssDNA-

MNP, and dsDNA-MNP samples were measured at 12–14 nM, and IgG-MNP and IgG-MNP-S1 (or S1*) samples were measured at 4 nM particle concentration.

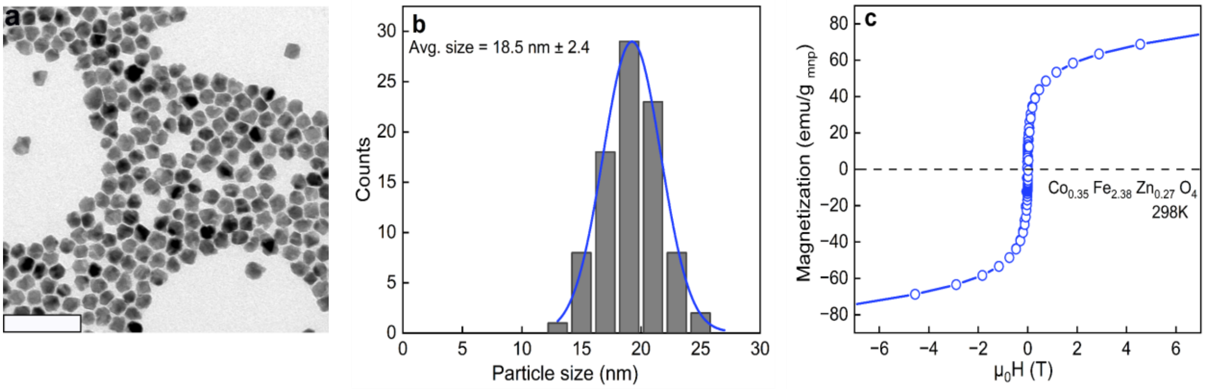


**Figure S1.** (a) Transmission electron microscopy (TEM) image of cubic-shaped nanoparticle used in this study. Scale bar is 100 nm. (b) Size distribution histogram as obtained from TEM image analysis, showing an average particle size of 18.5 ± 2.4 nm (n = 150). (c) Field-dependent magnetization hysteresis loops measured at 298 K and plotted as an average of three independent measurements.

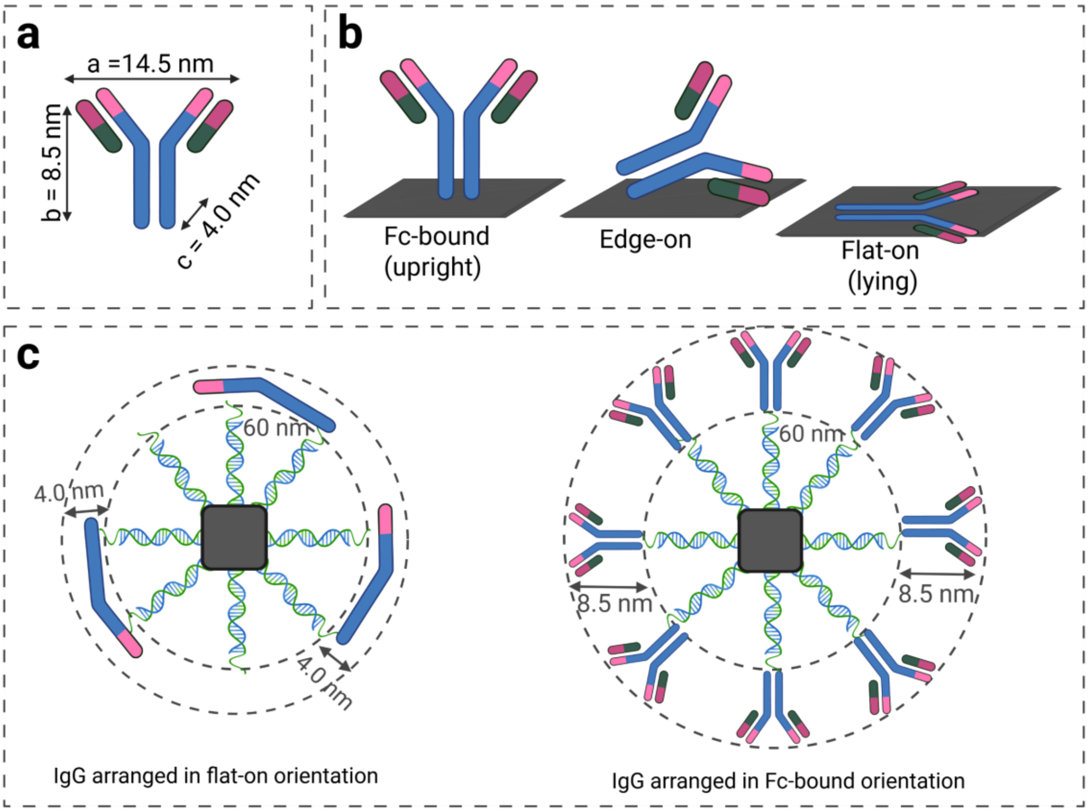


**Figure S2**. (a) Schematic illustration of monoclonal IgG antibody dimensions. (b) Three possible IgG orientations upon conjugation to MNP surface: Fc-bound, in which the Fc region of IgG anchors to the surface and both Fab arms project outwards; edge-on, in which the antibody stands on its narrow edge; and flat-on, in which the antibody lies with its largest face parallel to the MNP surface. (c) Schematic overview of IgG conjugation to ds-DNA-MNP (w/$N_3$) at low IgG loading (left) and high IgG loading (right). At low surface density, IgG adopts a flat-on orientation, with the thickness (dimension c) of 4.0 nm projecting outward from ds-DNA-MNP (w/$N_3$) shell, yielding an avearge size increase of ~ 8 nm (2×4.0). At high surface coverage, steric crowding on the dsDNA-MNP shell favors IgG binding in the Fc-bound orientation, with a height of 8.5 nm (dimension b) extending radially outward, yielding a size increase of ~ 17 nm (2×8.5). The inner circle represents the $D_h$ of dsDNA-MNP (w/$N_3$) with before IgG conjugation, and the outer circle represents the total $D_h$ after IgG conjugation.

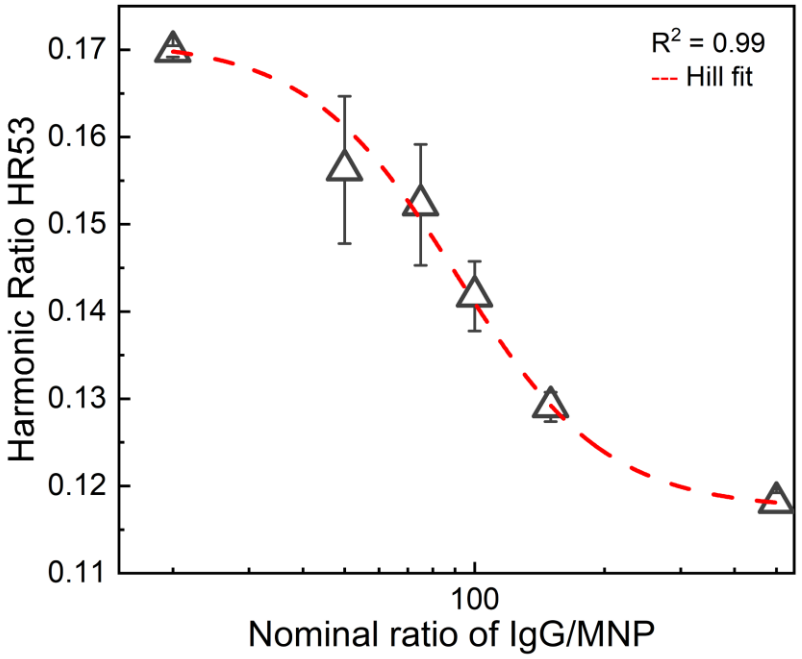


**Figure S3.** HR53 as a function of nominal IgG/MNP ratio prepared from dsDNA-MNP (with/$N_3$). As the ratio of IgG/MNP increases, particle hydrodynamic size increases, resulting in the decrease of HR53. Each data point represents the mean HR53 from three biological replicates, and error bars present the standard deviation between the replicates. The dashed red line indicates a Hill-like equation to the data with n = 2.5 ($R^2$ = 0.99). The $K_{1/2}$ value of ~ 91 indicates that HR53 reaches its half-minimum value at approximately 91 IgG/MNP.

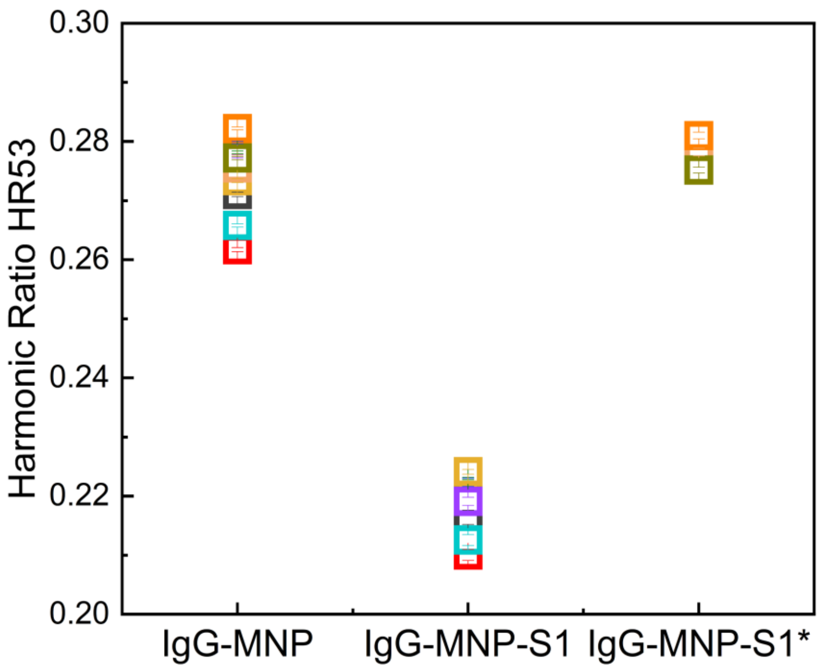


**Figure S4.** Individual HR53 measurements for the detection of SARS-Co-V-2 S1 protein using a fixed nominal IgG/MNP ratio of 150. This data is shown as box plots in Figure 3d of the main text. Ten samples of IgG/MNP were prepared, the target S1 subunit was added to seven of the IgG/MNP samples and S1* was added to the remaining three. Each coloured symbol represents an individual biological replicate. HR53 values decrease consistently across all IgG-MNP-S1 replicates compared to their paired IgG-MNP controls, while IgG-MNP-S1* values remain comparable to the control, confirming the specificity and reproducibility of the assay response.

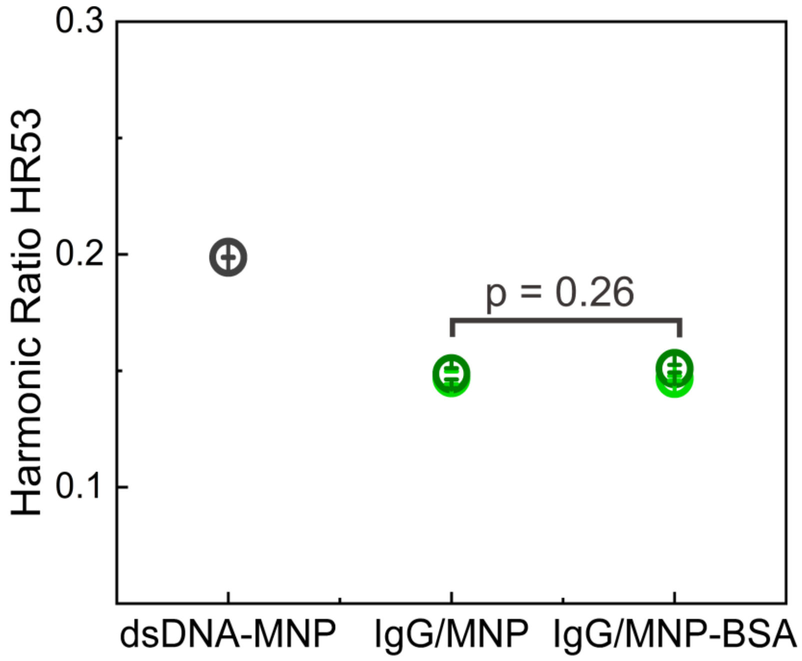


**Figure S5.** Harmonic ratio HR53 for dsDNA-MNP (w/N3), IgG-MNP conjugates prepared at 104 actual number of IgG/MNP, and IgG-MNP after addition of 0.5 % BSA. Two IgG/MNP samples were prepared at 4 nM particle concentration, excess IgG were removed with centrifugation and measured with MPS before adding BSA. The p-value of 0.26 was calculated based on the paired two-tailed t-test. The unchanged HR53 of IgG-MNP conjugates upon addition of BSA confirms the absence of non-specific BSA adsorption onto the IgG-MNP.

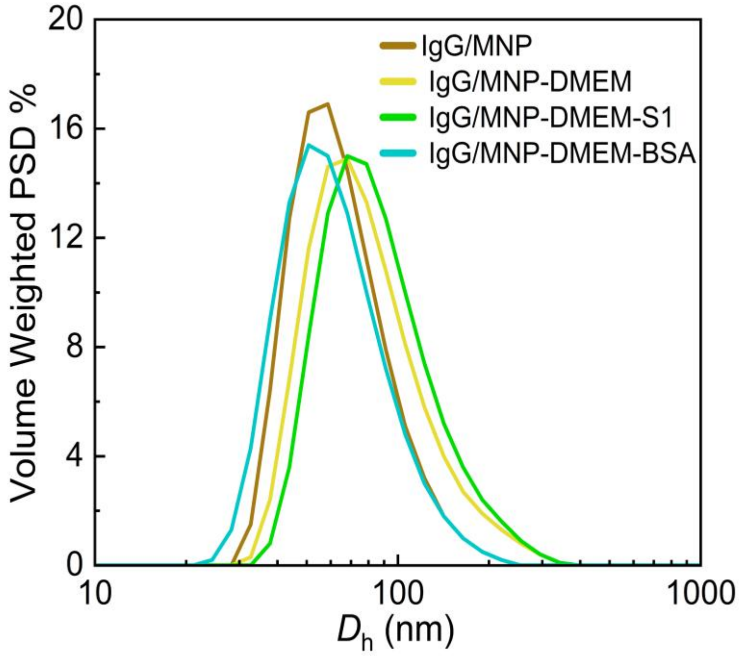


**Figure S6.** Volume-weighted particle size distribution of IgG-MNP (w/$N_3$) conjugates at each stage of assay in complex biological medium, as measured by DLS. S1 was added at a 3.6-fold molar excess relative to the IgG amount in the sample. The sequential shift in $D_h$ from IgG/MNP to IgG/MNP-DMEM and IgG/MNP-DMEM-S1 reflects the formation of protein corona followed by S1 binding. IgG/MNP-DMEM-BSA shows no shift relative to IgG/MNP, consistent with the absence of specific BSA binding.

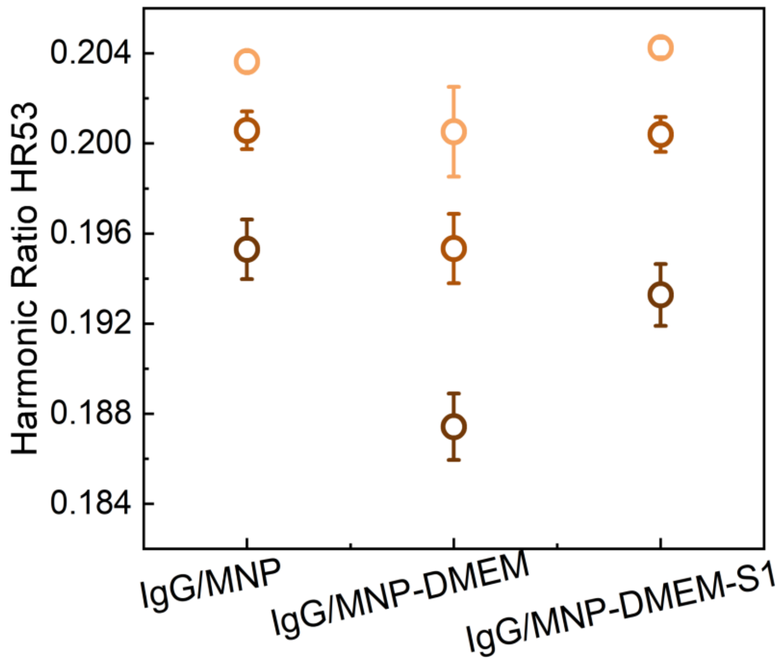


**Figure S7.** HR53 measured for IgG/MNP, IgG/MNP-DMEM, and IgG/MNP-DMEM-S1 at 1:1 molar ratio of IgG: S1. A decrease in HR53 was observed in IgG-MNP sample after DMEM addition, whereas no further signal drop is observed upon subsequent addition of S1. The absence of HR53 response to S1 at this ratio is likely due to insufficient amount of S1 to compete effectively with the off-target proteins adsorbed on the MNP surface. Individual data points of three biological replicates are shown.
